# Adsorption of Lysozyme Into a Charged Confining Pore

**DOI:** 10.1101/2021.07.11.451934

**Authors:** Daniel L. Z. Caetano, Ralf Metzler, Andrey G. Cherstvy, Sidney J. de Carvalho

## Abstract

Several applications arise from the confinement of proteins on surfaces since their stability and biological activity are enhanced. It is also known that the way a protein adsorbs on the surface is important for its biological function since its active sites should not be obstructed. In this study, the adsorption properties of hen egg-white Lysozyme, HEWL, into a negatively charged silica pore is examined employing a coarse-grained model and constant–pH Monte Carlo simulations. The role of electrostatic interactions is taken into account when including the Debye-HÃijckel potentials into the C*α* structure-based model. We evaluate the effects of pH, salt concentration, and pore radius on the protein preferential orientation and spatial distribution of its residues regarding the pore surface. By mapping the residues that stay closer to the pore surface, we find the increase of pH leads to orientational changes of the adsorbed protein when the solution pH gets closer to the HEWL isoelectric point. At these conditions, the pK_a_ shift of these important residues caused by the adsorption into the charged confining surface results in a HEWL charge distribution that stabilizes the adsorption in the observed protein orientation. We compare our observations to the results of pK_a_ shift for HEWL available in the literature and to some experimental data.

## 1 Introduction

The adsorption of proteins on confining surfaces, as routinely taking place in porous materials, has attracted considerable attention in recent years ^1,2^. This interest is due mainly to the broad applicability in biosensors, biocatalysts, and biomolecule delivery systems development ^3–6^. In these cases, the conditions of confinement lead to the protection of proteins against denaturation conditions and biological degradation, enhance their chemical and thermal stability, and, therefore, increase their biological activity. The study of native structure stability of proteins in nanopores has also provided relevant results for understanding the physical basis of protein folding ^7–9^. Furthermore, confined biomolecules are also used as models to study the biological transportation in biomembrane pores ^10,11^.

One of the determining factors for the applicability of confined proteins is their orientation regarding the surface and the consequent exposure of their active sites. The confinement effects on their native structure and the properties related to biological function, such as the local charge distribution, should be reduced ^12^. Experiments show that encapsulated proteins on charged surfaces exhibit an enhanced activity as compared to hydrophobic surfaces where this increase is highly dependent on the size and geometry of the cavity ^13–15^. In general, maximum adsorption is observed when the pH is close to the isoelectric point of the protein, and this trend is understood in purely electrostatic (ES) terms. At these conditions, the protein–protein repulsive interactions are minimal and the process is governed predominantly by the attractive protein–surface interactions ^16–19^.

Several computational studies have addressed the interactions between proteins and charged surfaces ^20,21^. Atomistic models were used to identify the preferential orientation and the most important residues for the adsorption. These studies focused, for instance, on Lysozyme and *α*-Chymotrypsin adsorption onto silica surfaces ^22,23^; Lysozyme, Cytochrome C, and Ribonuclease A adsorption onto self-assembled monolayers ^24–26^; and *β*-Lactoglobulin adsorption onto solid Au surfaces ^27^. Coarse-grained models have been adopted to investigate the salt and pH effects ^28–33^. An important aspect is the effect of surface ES potential on the degree of protonation of protein residues, the so-called charge regulation ^34–37^. In this case, a significant contribution to the binding free energy is observed at pH close to the isoelectric point of the protein and it is not noticed by constant–charge models ^37–39^. Despite of these efforts, molecular simulations of protein adsorption on charged confining surfaces are still scarce. ^40,41^.

In the current paper, we study the adsorption properties of hen egg-white Lysozyme (HEWL) into a negatively charged pore employing constant–pH Monte Carlo simulations. This protein is usually adopted as a model system to study protein adsorption onto porous materials ^42^. One of its functions is, for instance, the hydrolysis of the cell walls of Gram-positive bacteria ^43,44^. We adopt a coarse-grained model in which the ES interaction between the protein’s charged residues and between them and the surface is given by the Debye-Hückel potentials ^45,46^. The charges of titratable residues are affected not only by the solution conditions (salt concentration and pH), but also by the presence of the vicinal charged surface. We evaluate the influence of pH, salt concentration, and confinement radius on the protein–pore interactions and the distribution of charged residues relative to the pore surface. The effect of the confining–surface ES potential on the protonation degree of the residues is also analyzed, and its importance to the adsorption features is discussed.

The paper is organized as follows: In Sec. 2 we present the protein model employed and the details of our computer simulations. In Sec. 3 we examine the influence of salt concentration, pH, and confinement radius on protein–pore interactions and discuss the main findings. Finally, we conclude and outline some possible applications of our results in Sec. 4.

## 2 Model and Simulations

HEWL, see Fig. 1, is a small ellipsoidal enzyme constituted by a single chain with 129 amino-acid residues. Among its 129 residues, seven are aspartic acids (ASP), two are glutamic acids (GLU), three are tyrosines (TYR), one histidine (HIS), six are lysines (LYS), and eleven are arginines (ARG). The distribution of the residues results in an isoelectric point of about 1122. We model this protein according to the structure-based model (SBM), in which its residues are represented by single spheres centered at the C*α* position 49–54, and the non–ES energy is given by

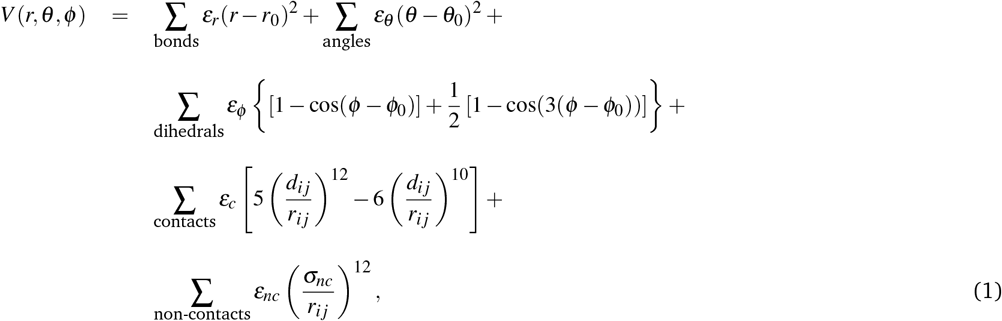

where *r*, *θ*, and *φ* are the distances between two consecutive residues, the angles formed by three consecutive residues, and the dihedral angles formed by four consecutive residues, respectively. The values of *r*_0_, *θ*_0_, and *φ*_0_ are obtained from the native structure of HEWL (PDB-ID 3WUN). The parameter *d_ij_* is the distance between residues *i* and *j* (*i < j* − 3) that are in contact in the native structure, according to the CSU (Contact of Structural Units) contact map ^55^. The parameters *∊_r_* = 100*∊_c_*, *∊_θ_* = 20*∊_c_* and *∊_φ_* = *∊_nc_* = *∊_c_* are defined as a function of the Lennard-Jones 10 − 12 parameter *∊_c_*. It determines the interaction energy between the residues in contact in the native structure and, accordingly, the transition temperature to the unfolded state. To set the value of *∊_c_*, we associate the unfolding–transition temperature obtained from the simulations, usually obtained from the peak of heat capacity, *C_v_* ^54^, with the melting temperature, *T_m_*, obtained experimentally. We calculate the heat capacities *C_v_* = (⟨*E*^2^⟨ − ⟨*E*⟨^2^)/*k_B_T*^2^ through the Weighted Histogram Analysis Method (WHAM), which calculates the density of states of the system at a given temperature *T*, using the energy histograms obatined from simulations at different constant temperatures. ^56,57^. Fig. 2 shows *C_v_* calculated from simulations for three different values of *∊_c_*, as well as *T_m_* provided in Ref. ^58^. Therefore, in this study, we define *∊_c_* = 0.63 kcal/mol to reproduce the melting temperature obtained experimentally. Finally, the parameter regarding the repulsive interactions between residues that do not belong to the native contact map is *σ_nc_* = 4.0 Å ^49,50^.

**Fig. 1.**
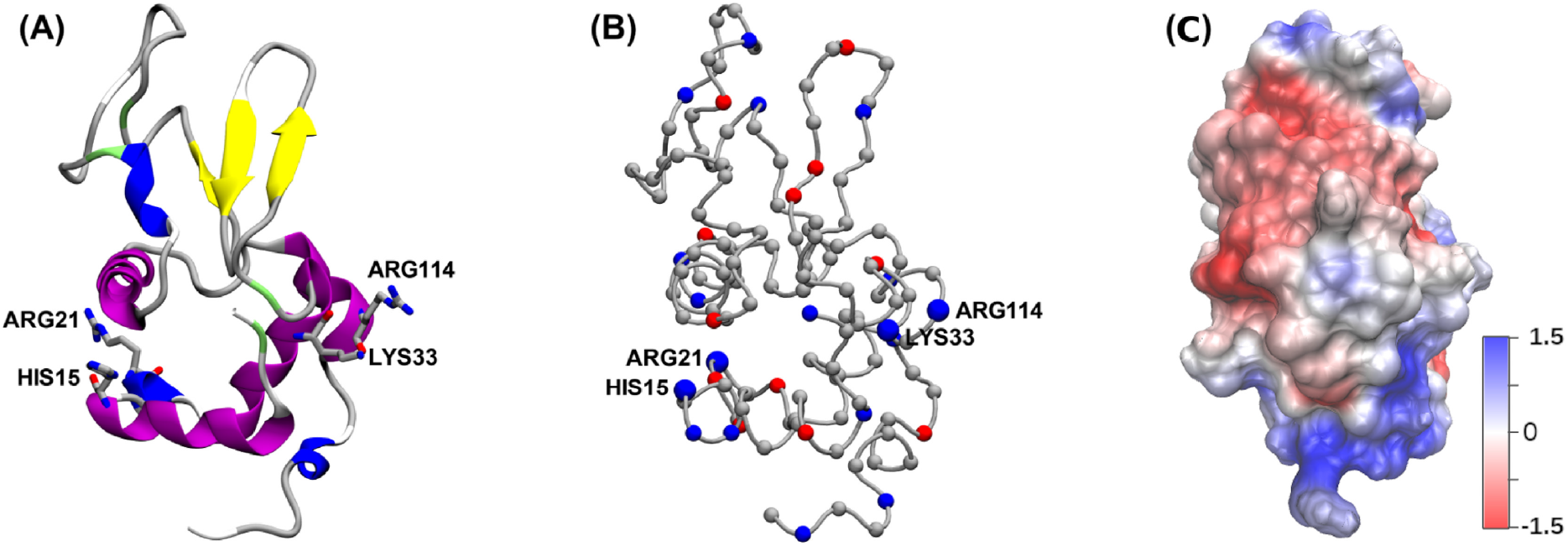
(A) Structure of hen egg-white Lysozyme (HEWL) protein built from PDB-ID 3WUN. The side chains of four important residues for HEWL– surface adsorption and orientation are shown. (B) C*α* representation of HEWL, where the basic residues (arginine, lysine, and histidine) are highlighted in blue, and the acidic ones (tyrosine, aspartic, and glutamic acids) are highlighted in red. (C) ES potential at the HEWL surface calculated using APBS Electrostatic Extension of VMD 47,48 considering the mean charges obtained from our simulation in the solution at the isoelectric point and salt concentration of 100 mM. The negative potentials are shown in red, while the positive ones are in blue. The unit of electrostatic potential is *k_B_T/e* (≈ 25 mV)

**Fig. 2.**
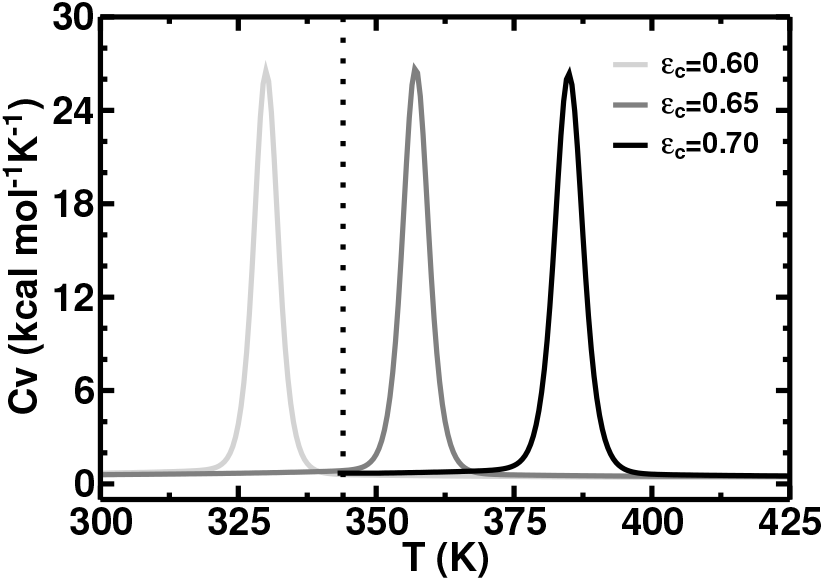
Heat capacities at constant volume, *C_v_*, in pH = 7.0 and salt concentration of 0.1 M for three different values of *∊_c_*: 0.60 kcal/mol (light gray line), 0.65 kcal/mol (gray line), and 0.70 kcal/mol (black line). We consider that the peak of *C_v_* as the melting temperature, *T_m_*. The vertical dotted line corresponds to the *T_m_* obtained experimentally in Ref. ^58^.

The ES interaction energy between any pair of charged residues is given by the screened Coulomb potential

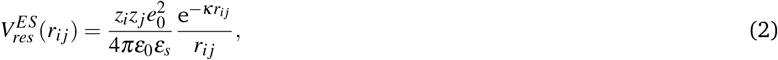

where *e*_0_ is the elementary charge, *z_i_* and *z_j_* are the respective valencies of residues *i* and *j*, *r_ij_* is the distance between them, *∊*_0_ is the permittivity of free space, *∊_s_* = 78.7 is the solvent dielectric constant, and *κ*^−1^ is the Debye screening length. This model has been successfully used to study the pH-dependent effects for protein folding and stability ^45,46,54^.

One single protein is confined within a cylindrical charged pore of radius *a* and surface charge density *σ_e_*, which is a function of pH of the solution. We obtain the values of *σ_e_* for the pH values adopted in this work from the experimental data for the silica nanoparticles presented in Ref. ^59^. The ES interaction energy between the charged residue *i* of HEWL inside the pore and the charged surface is obtained from the solution of the linear Poisson-Boltzmann equation given by ^60,61^

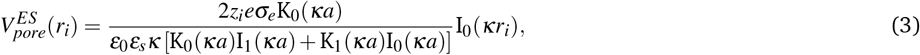

where *r_i_* is the radial distance between the residue *i* and the pore axis, and I_*n*_ and K_*n*_ are the modified Bessel functions of first and second kind, respectively. The Debye screening length is given by ^62^

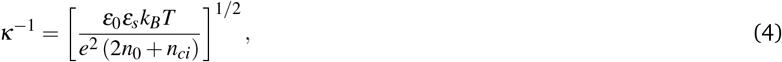

where *k_B_T* is the thermal energy, *n*_0_ is the number density of salt ions in the bulk of the solution, and *n_ci_* = (*σ_e_*2*πa*)*/*(*eπa*^2^) is the number density of the counterions to keep the system electroneutral.

To sample protein configurations, we carry out Metropolis Monte Carlo simulations in the canonical ensemble and used five kinds of movements: (1) translational displacement of a single residue randomly chosen, (2) radial displacement of the whole protein, (3) pivot rotation, (4) rotation of the whole protein, and (5) crankshaft movement ^63^. In each Monte Carlo step, we randomly choose one of these movements. To sample the protonation state of the residues, we also randomly choose 30% of configurations to try to change the protonation state of titratable residues. In these configurations, we randomly choose a titratable residue, and its protonation state is changed or not according to Metropolis criterion with the energy variation given by ^45,46,64^

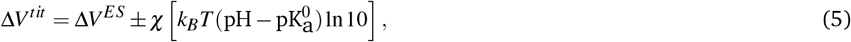

where *χ* = 1 or − 1 if the selected residue is basic or acidic, respectively. The positive and negative signs in Eq. (5) are used for protonation and deprotonation, respectively. The first term in Eq. (5) corresponds to the ES energy variation (Eqs. (2) and (3)) in the protonation/deprotonation. The second one is the compound model contribution, where the 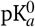 is the negative of the logarithm of the dissociation constant in the absence of electrostatic interactions with the other charged residues of the protein. The 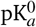 value of the titratable residues are 4.0 for aspartic acid, 4.5 for glutamic acid, 9.6 for tyrosine, 6.3 for histidine, 10.6 for lysine, and 12.0 for arginine ^46,65^. The cysteine is a titratable residue in its free form. HEWL has eight cysteine residues, but they form four disulfide bonds and cannot ionize. Therefore, in this work, they are considered to be neutral residues ^66^.

For each combination of pH, salt concentration and pore radius, we start the Monte Carlo simulations with the protein in its native state with all titratable residues charged. Then, the protein configurations are sampled using the movements described above and changing the state of protonation of the titratable residues. The equilibration process was carried out with 10^7^ Monte Carlo steps, and the average properties were calculated using 5 × 10^6^ statistically uncorrelated configurations of the protein.

## 3 Results and Discussions

We start our analysis by evaluating the dependence of the net charge of HEWL free in solution, *Q_p_*, on pH. Fig. 3 shows *Q_p_* obtained from simulations for salt concentrations of 0.001 (empty black circles) and the physiologically relevant salinity 0.1 M (filled black circles) in the absence of confinement. The computational results are in good agreement with the experimental ones (light gray stars) with the isoelectric point (pI ≈ 11) 67 unaffected by variations in the salt concentration *Cs*. The majority of the pK_a_ values obtained by simulations present no significant deviation from the experimental ones from Ref.^68^ for *Cs* = 0.1 M. For this comparison, we adopt the criterion of Goh and coauthors^69^ in which the deviation is considered significant only when the difference regarding the experimental values is higher than a 1.0 pK_a_ unit. The acidic residues GLU35, ASP66, and ASP87 of HEWL exhibit a deviation of 2.45, 1.36, and 1.43, respectively. Despite its high simplicity, the model adopted in this work is proved suitable to study acid-base equilibrium in proteins ^45,46,54^.

**Fig. 3.**
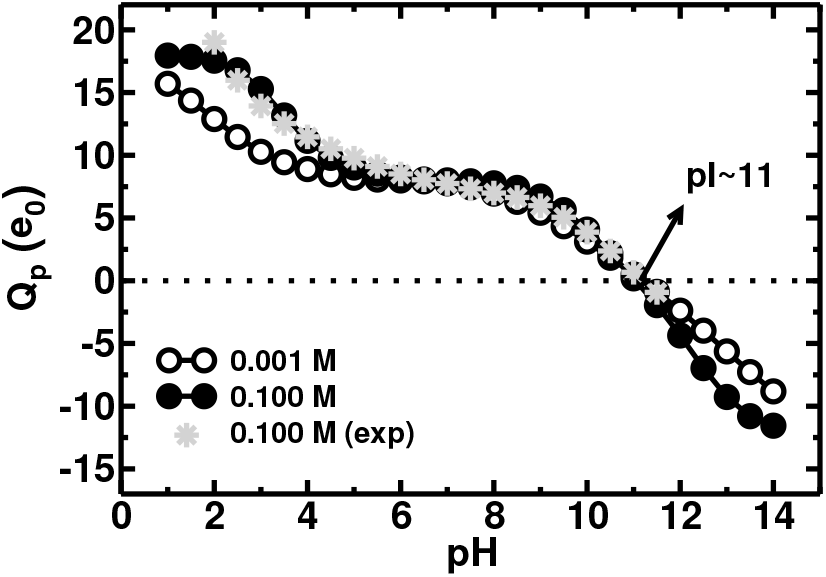
Net charge *Q_p_* (in units of elementary charges *e*_0_) of HEWL free in the solution as a function of pH, for salt concentrations of 0.001 (empty black circles) and 0.1 M (filled black circles). The experimental results (light gray stars) were obtained from Ref. ^67^.

To verify the conformational stability of the HEWL protein confined in the pore, we carried out simulations in regimes of low (0.001 M) and high (0.1 M) ionic strength and pore radii ranging from 40 to 800 Å (see Fig. S1 in Supplementary Information). The pH range considered was from 7 to 11 since, in these conditions, the protein charge and surface charge density result in stronger ES adsorption. For high salt concentration, the HEWL remains folded with approximately 90% of its native contacts formed (298 out of 330 contacts) and has the well-known mean radius of gyration *R_g_* ≈ 14 Å ^70^. In the weak–ES–screening regime corresponding to low salt concentrations, the protein undergoes a conformational transition in which the fraction of native contacts changes from about 0.9 to 0.55 at pore radius *a* = 400 Å. Perturbation on the HEWL secondary structure upon adsorption on silica nanoparticles was also observed in Ref. ^71^ from deconvolution of circular dichroism spectra.

Based on the analysis above, all of the following results under the conditions in which the protein has some flexibility but retains its native structure (*a* < 400 Å). Fig. 4 shows how pH, salt concentration, and pore radius affect the HEWL–surface binding energy, *E_B_*. For both salt concentrations (0.001 M in Fig. 4A and 0.1 M in Fig. 4B), *E_B_* presents a minimum value that becomes deeper and deeper and shifts slightly to higher pH with increasing pore radius. This increase in |*E_B_*| (in other words, it becomes more negative) occurs because the density of positive counterions, *n_ci_*, required to neutralize the negative silica surface, and so their contribution to the ES screening depends inversely on the pore radius *a* (see the definition of *n_ci_* in Eq. (4)). The displacement of the pH that corresponds to the strongest binding energy is partially related to the conditions which result in a maximal product between the protein charge and surface charge density and thus in maximal attraction of net-positively charged HEWL and negatively charged silica–surface.

**Fig. 4.**
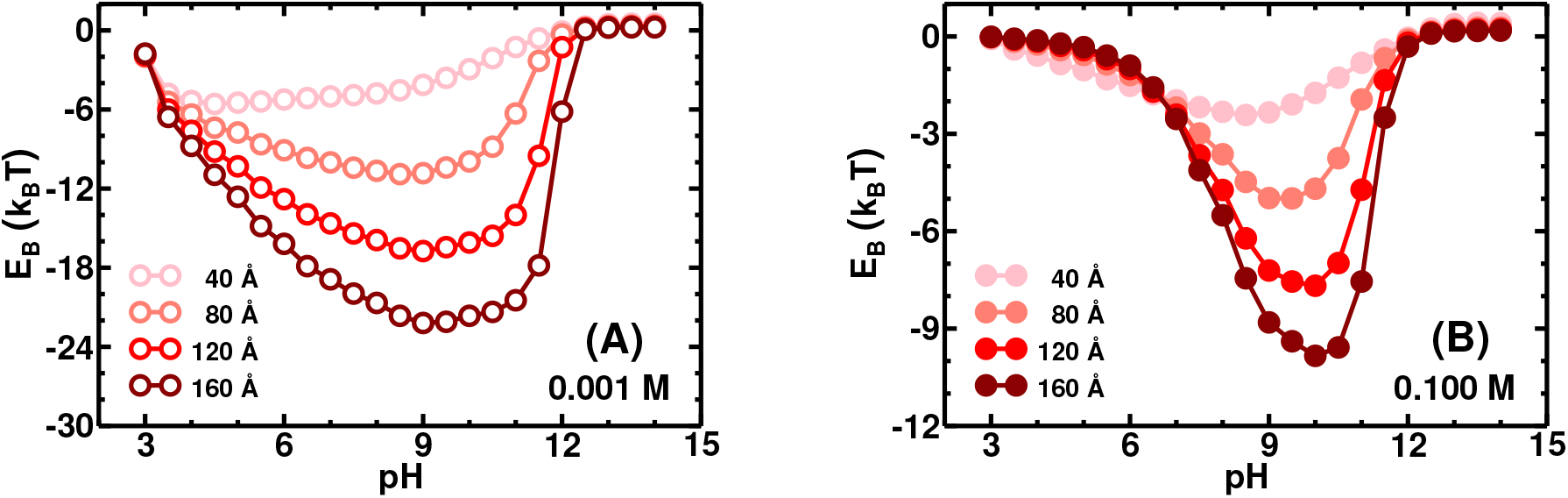
Mean binding energy, *E_B_*, of a single HEWL protein into a negatively charged pore as a function of pH for two salt concentrations: (A) 0.001 M - empty circles -, and (B) 0.1 M - filled circles. The pore radii are *a* = 40 Å (pink circles), 80 Å (light red circles), 120 Å (red circles) and 160 Å (dark red circles).

Another factor is the dependence of the directionality degree of the adsorbed protein on the pore radius, as will be seen later. When the binding is very directional, the adsorption is predominantly stabilized by the interaction of the pore surface with the specific positively charged region of the protein. We also see that increasing salt concentration not only decreases |*E_B_*|, but also diminishes the pH range in which HEWL is adsorbed. For pH < 6.5, the ES screening variation caused by increase salt concentration from 0.001 to 0.1 M leads from a state in which the protein is strongly adsorbed, and |*E_B_*| is about dozens k_*B*_Ts, to a weakly adsorbed state, where |*E_B_*| is about k_*B*_T. Once HEWL’s pI is approximately 11 and the silica’s point of zero charge is between pH 2 and 4^59^, we expected the protein is electrostatically attracted to the pore surface at pH between 2 and 11. However, Fig. 4A shows *E_B_* < 0 for pH regimes above this range. *E_B_* ≈ −6 μ 8 k_*B*_T at pH = 12 and low salt concentration, where *E_B_* fluctuations can be seen in Fig. S2A of Supplementary Information. This relatively high fluctuation of *E_B_* is predominantly due to protein charge fluctuations at this pH value, which corresponds to 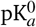 of arginine residues (see Figs. S2B and S2C in Supplementary Information). This fact suggests charge regulation is the possible driving force for adsorption at pH ≥ pI ^32,36,37^.

We evaluate the orientation of the adsorbed protein through the probability distribution *ρ*(*r*) of charged basic residues as a function of the distance *r* from the pore surface. Fig. 5 shows *ρ*(*r*) of residues HIS15 (yellow line), ARG21 (blue line), LYS33 (dark blue line), ARG61 (dark green line), ARG68 (dark yellow line), and ARG114 (green line) at low salt concentration (0.001 M) and pH = 7, 9, and 11. We focused here on the larger pore (*a* = 160 Å) to minimize the confinement effects. Since the radius of gyration of HEWL remains virtually constant in all cases presented in this study, the residues can be found at specific regions from the pore surface for each pH. Thus, we grouped the charged basic residues into three groups. Group 1 contains all residues that are more likely to locate at *r* ≤ 10 Å. Group 2 is composed of the ones that are more likely to find at 10 < *r* ≤ 20 Å, and the ones that are more likely to locate at *r* > 20 Å belong to Group 3. Table 1 shows the details of all charged basic residues which belong to each group for the pHs shown in Fig. 5.

**Fig. 5.**
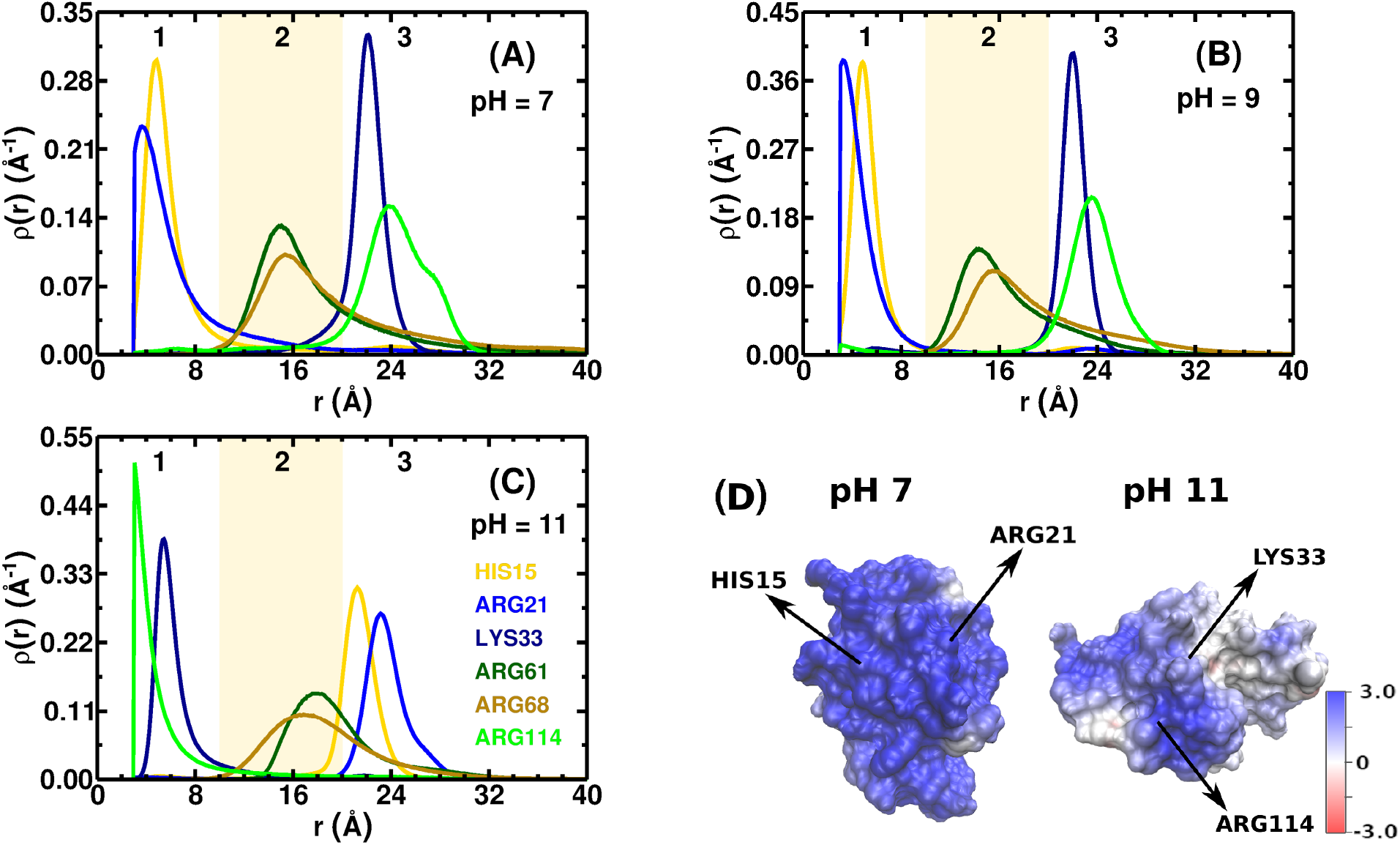
Probability distribution of the residues HIS15 (yellow lines), ARG21 (blue lines), LYS33 (dark blue lines), ARG61 (dark green lines), ARG68 (dark yellow lines), and ARG114 (green lines) as a function of the distance *r* from the pore surface for three pHs: (A) pH = 7, (B) pH = 9, and (C) pH = 11. The pore radius is *a* = 160 Å and the salt concentration is 0.001 M. Regions 1, 2 and 3 correspond to the groups of basic residues which are more likely located at *r* 10 Å (Group 1), 10 Å < *r* 20 Å (Group 2), and *r* > 20 Å (Group 3), respectively, as detailed in Table 1. (D) ES potential at the HEWL surface calculated by means of APBS Electrostatic Extension of VMD ^47,48^ considering the mean charges obtained from our simulation at pH = 7 and 11 and salt concentration of 0.001 M. The negative potentials are shown in red and positive ones are in blue. The unit of electrostatic potential is *k_B_T/e* (≈ 25 mV).

**Table 1.** List of basic residues which are more likely located at *r* ≤ 10 Å (Group 1), 10 Å < *r* ≤ 20 Å (Group 2), and *r* > 20 Å (Group 3) from the pore surface. The pore radius is *a* = 160 Å and the salt concentration is 0.001 M.

| pH | Group 1 | Group 2 | Group 3 |
| --- | --- | --- | --- |
| 7 | LYS13, ARG14, HIS15<br>ARG21, ARG73, LYS96<br>LYS97 | ARG5, ARG61, ARG68<br>ARG112, LYS116, ARG125<br>ARG128 | LYS1, LYS33, ARG45<br>ARG114 |
| 9 | LYS13, ARG14, HIS15<br>ARG21, ARG73, LYS96<br>LYS97, ARG128 | ARG5, ARG61, ARG68<br>ARG112, LYS116, ARG125 | LYS1, LYS33, ARG45<br>ARG114 |
| 11 | LYS1, ARG5, LYS33<br>ARG45, ARG112, ARG114<br>LYS116, ARG125, ARG128 | LYS13, ARG14, ARG61<br>ARG68 | HIS15, ARG21, ARG73<br>LYS96, LYS97 |

Another feature we observe in Fig. 5 is that the increasing pH value promotes an orientational change of HEWL. Although the protein is adsorbed on the pore surface for the three pH values shown in Fig. 5 (see also the dark red circles in the Fig. 4A), we noticed an inversion between the residues belonging to Groups 1 and 3 caused by the increasing of pH towards HEWL’s pI (pI ∼ 11). Fig. 5D shows the side of the protein that is in contact with the pore in the two observed orientations at pH = 7 and 11 and *C_s_* = 0.001 M. The arrows point out the location of the two closest residues to the pore surface. The protein surface is colored according to the ES potential and highlights the positive region (in blue) where these residues are located. Fig. 6 shows the probability distribution *ρ*(*r*) of the six basic residues considered in Fig. 5, under the same conditions, except at pH = 10. In this case, *ρ*(*r*) is bimodal since the protein changes its orientation with respect to the surface during the simulation. Snapshots of these two orientational states are shown in Figs. 6B and 6C.

**Fig. 6.**
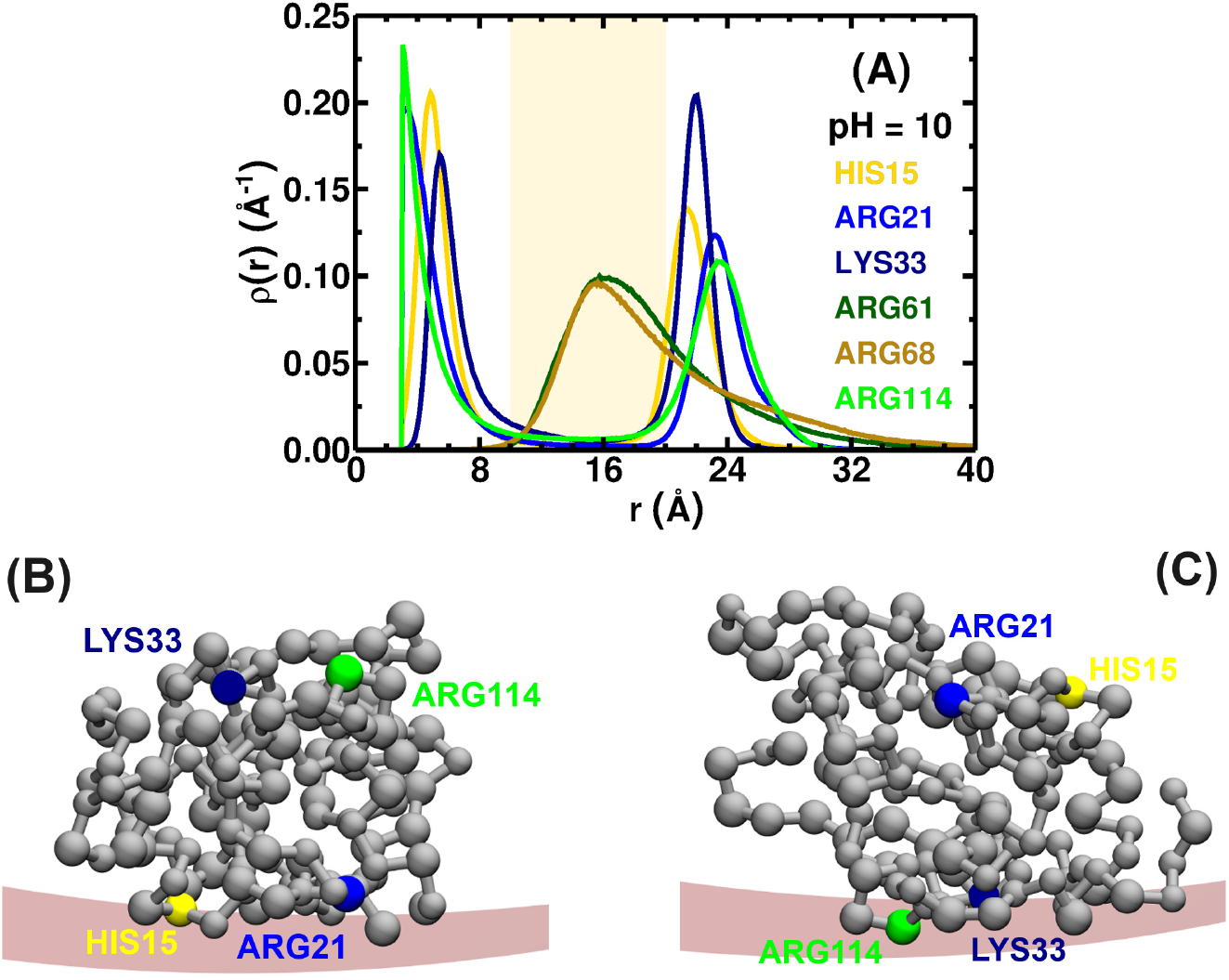
Probability distribution of the residues HIS15 (yellow line), ARG21 (blue line), LYS33 (dark blue line), ARG61 (dark green line), ARG68 (dark yellow line), and ARG114 (green line) as a function of the distance *r* from the pore surface. The pore radius is *a* = 160 Å, the salt concentration is 0.001 M and pH = 10. The snapshots shown in (B) and (C) correspond to two orientational states of HEWL with the residues HIS15, ARG21, LYS33, and ARG114 highlighted in yellow, blue, dark blue and green, respectively.

Experimental investigations have demonstrated that the adsorption of HEWL is such that its short axis is perpendicular to the surface with N, C-terminus faces playing an important role (side-on orientation) ^20,72–74^. In a series of studies using detailed atomistic Molecular Dynamics simulations, Kubiak-Ossowska and coauthors have focused on the adsorption of HEWL on flat charged silica surfaces at pH = 7^40,75–77^. They have found that ES interactions play a key role in guiding HEWL to the most favorable orientation with N, C-terminal face against the silica surface. Besides that, they have listed a set of residues – such as LYS1, ARG5, LYS13, ARG14, ARG125, and ARG128 – that are important for protein–surface anchoring. Yu and Zhou have studied the curvature effect of silica nanoparticles on the Lysozyme orientation and conformation by Molecular Dynamics simulations of a mesoscopic coarse-grained model at neutral pH ^31^. They have verified that the number of binding sites increases with increasing nanoparticle curvature and decreasing salt concentration. Four of the seven residues belonging to Group 1 at pH = 7 in our results are the same as those mentioned by these works. This difference can be attributed to the charge regulation effects in our work, whereas the studies cited above ^31,40,75–77^ consider the residues with fixed pH independent charge. Recently, Boubeta and coauthors have proposed a theoretical framework to study the interaction between the Lysozyme and charged flat surfaces at pH = pI, taking into account charge–regulation effects ^32^. They have found that LYS1, ARG5, LYS33, ARG114, ARG125 and ARG128 are much closer to the surface and likely more important for the protein-surface interaction. Our findings (see the bottom line in Table 1 and Fig. 5C) are not only in agreement with these results, but also with the experimental data from Dismer and colleagues who developed an effective lysine–labeling technique to identify possible binding orientations of Lysozyme on surfaces ^74^. A change of the protein orientation at pH > 10 is also observed experimentally by Kubiak-Ossowska and coauthors but attributed to the dominance of hydrophobic forces ^22^. In our case, we notice an orientational change even considering only electrostatic interactions.

Figures 7A-7C show the salt concentration effects on the probability distribution *ρ*(*r*) of one residue of each Group defined above when the protein is within a pore of radius *a* = 160 Å at pH = 9. The increase in the salt concentration results in a decrease in the probability of ARG21 (Group 1) being near the pore surface. The same effect occurs for LYS33 (Group 3) be far from the pore surface. Furthermore, *ρ*(*r*) presents a second peak for the residues belonging to Groups 1 and 3. The higher peak of the residue ARG21 (Group 1) corresponds approximately to the smaller peak of the residue LYS33 (Group 3) and vice versa. The residue ARG61, which belongs to Group 2, has a single peak at intermediary distances from the pore surface. As HEWL does not lose its native structure and its gyration radius remains constant, this behavior confirms the existence of two orientational states of the protein. Although the HEWL is adsorbed predominantly with residues from Group 1 in contact with the pore surface, the probability of the residues from Group 3 to be close to the surface is non-negligible as well. Nevertheless, the directionality of the adsorbed protein is getting lost with the increase of ionic strength due to screening of protein–surface ES interactions.

**Fig. 7.**
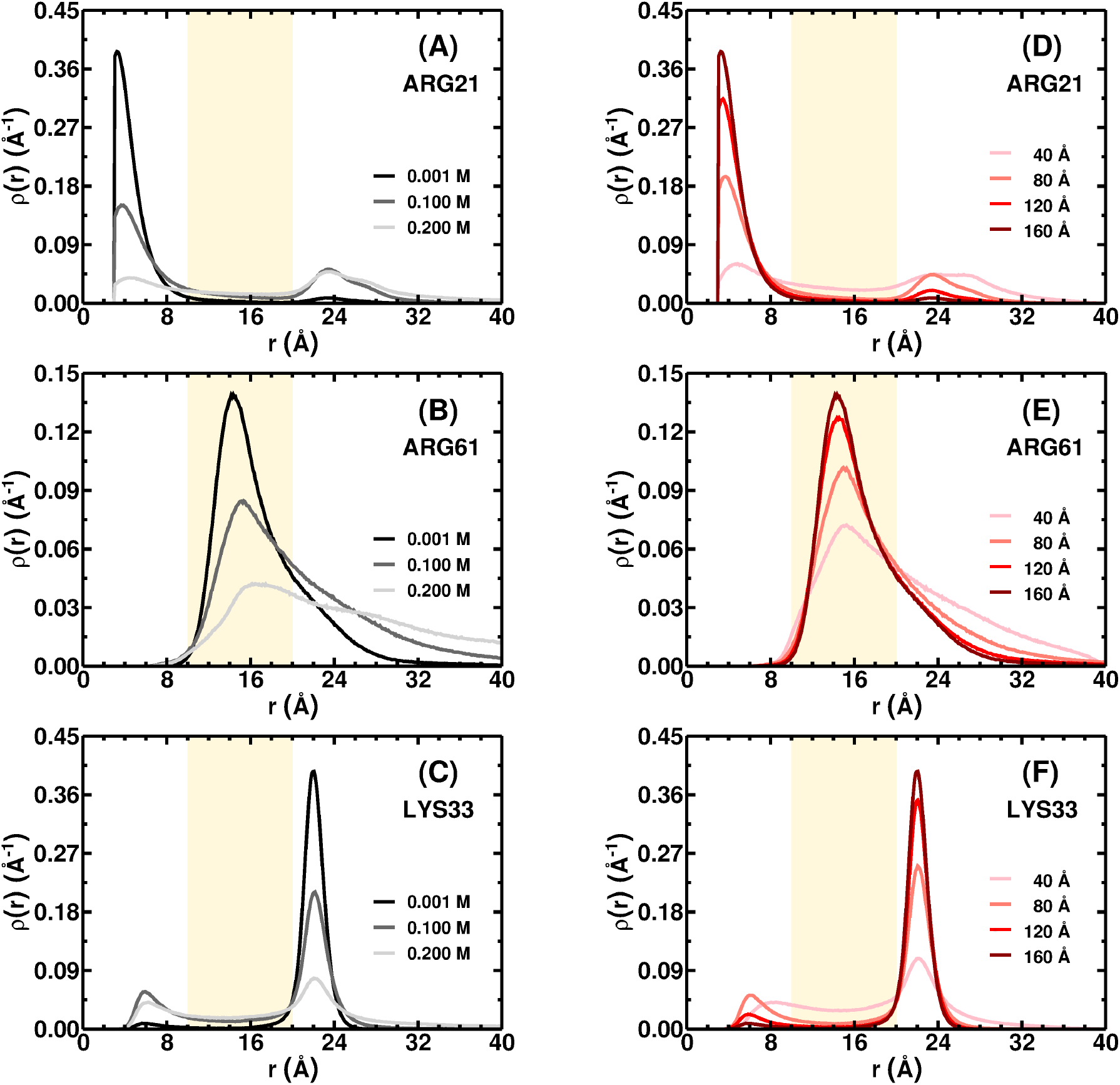
Probability distribution of the residues ARG21 (graphs A and D), ARG61 (graphs B and E), and LYS33 (graphs C and F) as a function of the distance *r* from the pore surface. In the left column (graphs A, B, and C), we set the pore radius in *a* = 160 Å and vary the salt concentration: 0.001 M (black lines), 0.1 M (gray lines), and 0.200 M (light gray lines). In the right column (graphs D, E, and F), we set the salt concentration at 0.001 M and vary the pore radius: *a* = 40 Å (pink lines), *a* = 80 Å (light red lines), *a* = 120 Å (red lines), and *a* = 160 Å (dark red lines). In all graphs, we have pH = 9.

The pore radius effect can be seen in Figs. 7D-7F, which show the probability distribution *ρ*(*r*) of residues ARG21, ARG61, and LYS33 for pore radius *a* = 40 Å (pink line), 80 Å (light red line), 120 Å (red line), and 160 Å (dark red line) at pH = 9 and 0.001 M of salinity. The reduction of the pore radius decreases the ES attraction between the protein and the surface. As already shown, the ES binding energy of adsorbed HEWL decreases (becomes less negative) with decreasing pore radius (see Fig. 4A and the discussion after it). The probabilities of the two orientations states approach each other with decreasing pore radius. In the smallest pore radius, *a* = 40 Å, we observed non-negligible probabilities of the residues being at intermediate positions between two peaks, indicating the loss of directionality of the adsorbed protein.

We also verify how the adsorption of HEWL on a charged confining surface modify the pK_a_ of protein residues. To achieve this goal, the protonation degree *α* of each ionizable residue was calculated at several pH conditions for the free protein in solution and when confined in a pore. The pK_a_ was then obtained by fitting the individual titration curves using the Hill equation ^78–81^

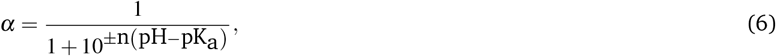

where the plus and minus signs are used for basic and acidic residues, respectively, and *n* is the Hill coefficient. This parameter provides information about the effect of the correlation between the residues in the course of the protonation. A Hill coefficient *n* > 1 means that the protonation is cooperative, i.e., the protonation of one residue is favored by the protonation of other ones, whereas if *n* < 1, the proton binding is anti-cooperative. In the case of *n* = 1, the residues behave independently of the others regarding the propensity towards protonation. In this work, for the HEWL free in solution, *n* < 1 for all titratable residues. As the protein is within the pore, *n* > 1 for the residues from Group 1 showed in Table 1 for pH = 11 (see Tables S1 and S2 in Supplementary Information). At this pH regime, the LYS and ARG residues are more susceptible to the charge–regulation effects since 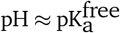, where the label “free” refers to the free protein in the solution. The protonation of one of these residues can cause variations in the strength and characteristics of adsorption that increases the ES potential effect on another one. The Hill coefficient approaches 1 with increasing ionic strength and decreasing pore radius due to the decreasing ES potential.

Table 2 shows the 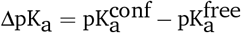 of HEWL residues at *C_s_* = 0.001 M that have |ΔpKa| > 1.0^45,46^, where the label “conf” refers to the confined protein. The action of the ES potential of the pore surface on the ionizable residues changes their probability of protonation. Furthermore, we verified that the reduction of the pore radius decreases |*E_B_*| and, consequently, the modulus of the ES potential (see Fig. 4A). As we can see in Table 2, lysine and histidine residues have a more pronounced pK_a_ shift for *a* = 160 Å, while arginine residues show a greater change in their pK_a_ in the smallest pore radius (*a* = 40 Å). The increase of the ES screening at 0.1 M of salt led to |ΔpKa| < 1.0 for all titratable residues and confinement conditions studied in this work.

**Table 2.** The pKa shift 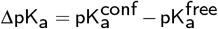 of basic residues from HEWL at low salt concentration (0.001 M).

| Residues | $\Delta pK_a$ | | | |
| --- | --- | --- | --- | --- |
| | $a = 160 \text{ \AA}$ | $a = 120 \text{ \AA}$ | $a = 80 \text{ \AA}$ | $a = 40 \text{ \AA}$ |
| LYS1 | $1.01 \pm 0.02$ | $0.73 \pm 0.03$ | $0.32 \pm 0.02$ | $-0.04 \pm 0.02$ |
| ARG5 | $-0.95 \pm 0.02$ | $-1.04 \pm 0.02$ | $-1.14 \pm 0.02$ | $-1.25 \pm 0.02$ |
| ARG14 | $-0.95 \pm 0.02$ | $-1.03 \pm 0.02$ | $-1.12 \pm 0.02$ | $-1.20 \pm 0.02$ |
| HIS15 | $1.56 \pm 0.02$ | $1.34 \pm 0.02$ | $1.18 \pm 0.02$ | $1.14 \pm 0.02$ |
| ARG21 | $-1.85 \pm 0.02$ | $-1.95 \pm 0.02$ | $-2.09 \pm 0.02$ | $-2.30 \pm 0.02$ |
| LYS33 | $1.15 \pm 0.03$ | $0.86 \pm 0.03$ | $0.43 \pm 0.02$ | $0.04 \pm 0.02$ |
| ARG45 | $-1.20 \pm 0.02$ | $-1.30 \pm 0.02$ | $-1.40 \pm 0.02$ | $-1.54 \pm 0.02$ |
| ARG61 | $-0.99 \pm 0.02$ | $-1.08 \pm 0.02$ | $-1.16 \pm 0.02$ | $-1.25 \pm 0.02$ |
| LYS116 | $1.55 \pm 0.06$ | $1.18 \pm 0.05$ | $0.82 \pm 0.03$ | $0.57 \pm 0.03$ |

In order to understand the inverse behavior of arginine residues with respect to change of their pK_a_ with the pore radius, Fig. 8A shows the degree of protonation *α*_ARG21_ of ARG21 residue for the HEWL free in the solution (purple color) and within the pore with radius *a* = 40 Å and 160 Å (pink and dark red colors, respectively) at *C_s_* = 0.001 M. The protonation degree of ARG21, at pH ≥ 11 is as larger as the ES potential *ϕ*_ARG21_ acting on this residue is negative (see Fig. 8B). For pH > 12, the |*ϕ*_ARG21_| regarding the free protein in solution becomes larger, promoting the protonation of ARG21. The sensitivity of the protonation state to the action of the electrical potential from the pore surface is larger for 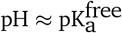. In the case of ARG21, 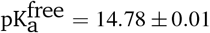 and the inversion of the ES potential behaviour in which *ϕ*_ARG21_ regarding the protein free in solution becomes more negative occurs for pH > 11. In the case of LYS116 (see Figs. 8C and 8D), which presents the highest positive ΔpKa, this inversion of behaviour occurs at pH ≈ 12.5 and 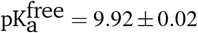. As a consequence, the protonation process of LYS116 is favored when the HEWL is confined to a pore with radius *a* = 160 Å.

**Fig. 8.**
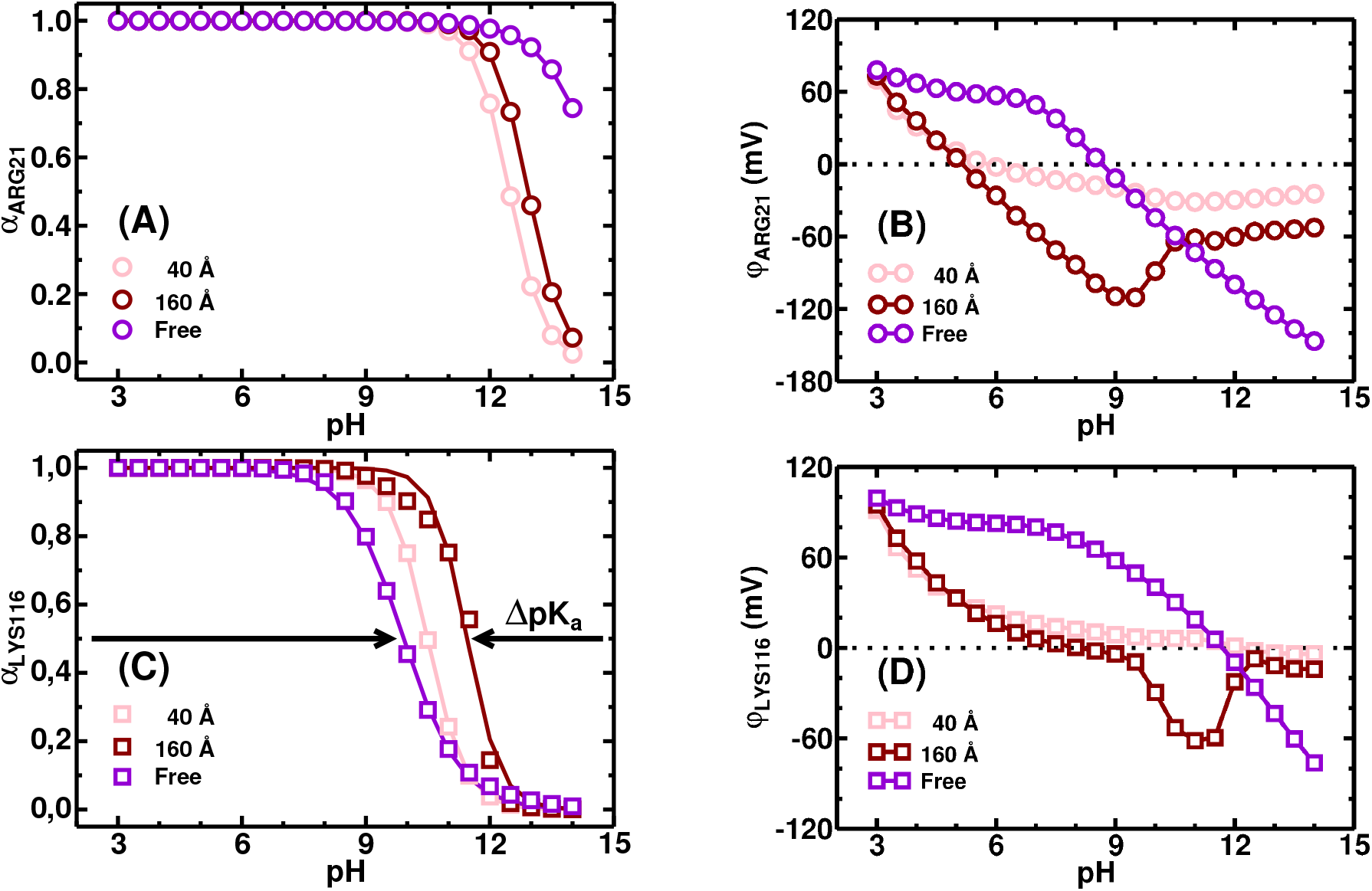
Protonation degree, *α*, of the residues ARG21 (A) and LYS116 (C) in the presence (pink and dark red colors) and absence (purple color) of confinement. Empty circles and squares are the results from Monte Carlo simulation for the residues ARG21 and LYS116, respectively, whereas the lines represent the fits using the Hill equation (Eq. (6)). ES potential at the residues ARG21 (B) and LYS116 (D) in the presence (pink and dark red symbols) and absence (purple symbols) of confinement. In all cases, *C_s_* = 0.001 M.

This pK_a_ shift causes a change in the protein charge distribution and its net charge, as shown in Table 3 for pore radius *a* = 160 Å, low ionic strength (0.001 M), and pH = 7 and 11. At pH = 7, far from the HEWL isoelectric point, the changes of the ES potential on the protein surface in contact with the pore surface are small; the same is observed for the protein net charge. Conversely, close to the isoelectric point of the protein, at pH = 11, the net charge grows about six times. The ES potential on the charge patch that contains the residues LYS33 and ARG114 also changes in this same proportion. This local induced charge density stabilizes the protein adsorption with the features shown in Fig. 5C.

**Table 3.**
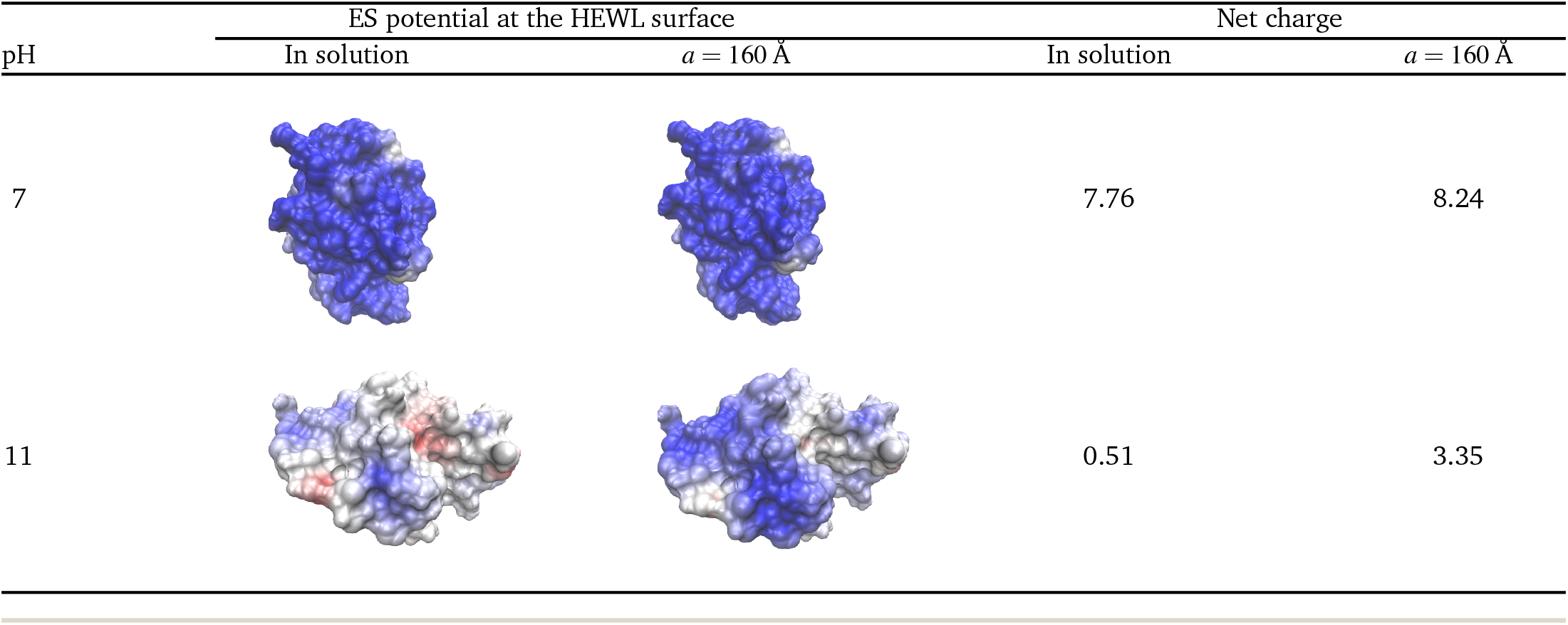
The net charge and ES potential at the HEWL surface calculated by means of APBS Electrostatic Extension of VMD ^47,48^ considering the mean charges obtained from our simulation at pH = 7 and 11 and *C_s_* = 0.001 M. The negative potentials are shown in red and positive ones are in blue. The color scale is the same as in Fig. 5

| pH | ES potential at the HEWL surface |  | Net charge |  |
| --- | --- | --- | --- | --- |
| | In solution | $a = 160 \text{ \AA}$ | In solution | $a = 160 \text{ \AA}$ |
| 7 |  |  | 7.76 | 8.24 |
| 11 |  |  | 0.51 | 3.35 |

This is consistent with results by Narambuena and coauthors who have studied the adsorption of lysozyme on weak polyacid hidrogel films using a molecular theory ^82^. They have verified that lysozyme adsorption occurs even under conditions where pH > pI due to charge–regulation effects. For example, they showed that, at pH = 11.3, the charge of the protein is approximately −2*e*_0_ when it is in bulk solution and, when the Lysozyme is adsorbed, its charge increases up to approximately +7*e*_0_.

## 4 Conclusions

In this study, we investigated the adsorption of hen egg-white lysozyme (HEWL) onto a negatively charged confining silica pore by means of constant-pH Monte Carlo simulations. We found that, except for the smallest pore radius, the protein-pore binding energy presents a minimum near HEWL’s isoelectric point, suggesting that charge patches and the charge regulation process play an important role for adsorption, see Fig. 4. We identified that the HEWL adsorbs in two orientations dependent on pH. At neutral pH, the HIS15 and ARG21 are the residues closer to the pore surface. Increasing the pH towards the pI, the orientation changes with the residue ARG114 having an important role for the adsorption stability, see Fig. 5 and 6. In addition, as shown in Tables 1–3, we group the residues most likely to be close to the pore surface and found the protonation degree of these residues is significantly affected by the protein– pore ES interactions, thereby favoring protein adsorption. Extensions of the current study include the adsorption at different protein concentrations, which will enable us to evaluate the cooperative effects caused by protein–protein interactions and, accordingly, the factors determining the maximum loading inside the pore. As pointed out by Moerz and Huber 19, the presence of charged patches and charge regulation effects can be important for the adsorption of Cytochrome C and Myoglobin into SBA-15 mesoporous silica materials, where the maximum pore loading is observed at pH values below the isoelectric point.

The results presented in this study provide valuable information on the adsorption of proteins on charged materials and can be applied to a wider range of problems. Even adopting a simplified model for the protein, we obtained a good agreement with results from more detailed models and from experiments. An immediate application of our results includes the problems of protein adsorption in porous media. The maximum load within the charged pores is strongly dependent on protein–protein interaction and conditions on its net charge. Besides, the detailed protein charge distribution determines the orientarion regarding the pore surface and of each protein with each other. These protein properties should be known when the protein is inside a pore and, as is shown by the results presented in this work, the changes of the protein properties regarding the protein free in the solution can be significant. This charge–regulation mechanism promotes not only a significant protein net charge at pH values close to the isoelectric point, that enhance the HEWL–silica adsorption, but also the distribution and charge density of charge patches on the protein surface, that determines the protein orientation regarding the pore surface. Here, returning to the idea of protein and drug delivery into a biological cell by nanoparticles of mesoporous silica, we emphasize that the shifts of pK_a_ for the protein residues found for HEWL in small pores should be tested for other proteins and ubiquitously used drug molecules. The shifts of pKa for particular residues, depending on the pore dimensions and the pH level of the solution, can potentially present a mechanism for release of the drug molecules inside the target cancer cells (which typically feature smaller pH values than healthy tissues/cells).

Another possible application includes the adsorption of lytic peptides onto anionic lipid membrane. Recently, it was shown that the lipid ES potential and the solution pH affect the degree of protonation of peptides residues and modulates the affinity to the membrane ^83^. It is known that at a given critical concentration, these peptides form pores in the bilayer that leads to the cell–death of bacteria. Therefore, to know the peptides’ charge distribution inside these charged confined enviroments should be essential to evaluate the driving forces and mechanisms of pore formation.

## Conflicts of interest

There are no conflicts to declare.

## Acknowledgements

All simulations were performed by resources supplied by the Center for Scientific Computing (NCC/GridUNESP) of the Sao Paulo State University (UNESP). Daniel L. Z. Caetano thanks the Sao Paulo Research Foundation (FAPESP, Grant No. 2013/08293-7 and 2019/19662-0) and the Coordenação de Aperfeiçoamento de Pessoal de Nível Superior, Brasil (CAPES), Finance Code 001 for the financial support. Ralf Metzler acknowledges support from the Foundation for Polish Science (FNP) within an Alexander von Humboldt Honorary Polish Research Scholarship. Sidney J. de Carvalho thanks to Sao Paulo Research Foundation (FAPESP, Process Grant No 2018/01841-2) for the financial support.

## Notes

### Competing Interest Statement

The authors have declared no competing interest.

